# Megakaryocyte Progenitor Cell Function is Enhanced Upon Aging Despite the Functional Decline of Aged Hematopoietic Stem Cells

**DOI:** 10.1101/2021.01.28.428507

**Authors:** Donna M. Poscablo, Atesh K. Worthington, Stephanie Smith-Berdan, E. Camilla Forsberg

## Abstract

Age-related morbidity is associated with a decline in hematopoietic stem cell (HSC) function, but the mechanisms of HSC aging remain unclear. We performed heterochronic HSC transplants followed by quantitative analysis of cell reconstitution. While young HSCs outperformed old HSCs in young recipients, young HSCs unexpectedly failed to outcompete the old HSCs of aged recipients. Interestingly, despite substantial enrichment of megakaryocyte progenitors (MkPs) in old mice *in situ* and reported platelet (Plt) priming with age, transplanted old HSCs were deficient in reconstitution of all lineages, including MkPs and Plts. We therefore performed functional analysis of young and old MkPs. Surprisingly, old MkPs displayed unmistakably greater regenerative capacity compared to young MkPs. Transcriptome analysis revealed putative molecular regulators of old MkP expansion. Collectively, these data demonstrated that aging affects HSCs and megakaryopoiesis in fundamentally different ways: whereas old HSCs functionally decline, MkPs gain expansion capacity upon aging.

**HIGHLIGHTS:**

- Frequencies and total cell numbers of HSCs and MkPs were increased upon aging
- Reconstitution deficit by old HSCs was observed by chimerism and absolute cell numbers
- Young HSCs did not have competitive advantage over old HSCs in aged recipient mice
- Old MkPs display remarkable capacity to engraft, expand, and reconstitute platelets
- Aging is associated with changes in MkP genome-wide expression signatures

## INTRODUCTION

The elderly population is affected by numerous hematological abnormalities, including decreasing capacity to mount an immune response, increasing incidences of thrombotic cardiovascular disorders, and dramatically increased incidence of myelogenous diseases. Dysfunctions during chronological aging is paralleled by the declining functionality of hematopoietic stem cells (HSCs), leading to conclusions that the age-related pathologies of the blood manifests within the HSC compartment (Elias *et al.*, 2017). HSCs self-renew and differentiate, giving rise to progenitor cells that are committed to either the myeloid or lymphoid lineage throughout the lifespan. However, there is convincing evidence that HSCs lose the full extent of these properties during aging. Evaluation of aged HSCs in transplantation studies has demonstrated reduced regenerative potential compared to young HSCs. Aged HSCs seem to be less effective at homing and engrafting in the host, suggesting that alterations to HSCs during aging persist (become intrinsic) upon transplantation (Morrison *et al.*, 1996; Sudo *et al.*, 2000; Rossi *et al.*, 2005; Dykstra *et al.*, 2011). In those experiments, HSC multipotency has been traditionally defined by qualitative measurements of GM, B, and T cell donor-to-host chimerism; more quantitative measurements and direct tracking of erythroid cells and platelets (Plt) from hematopoietic stem and progenitor cells (HSPCs) were developed relatively recently to rectify limitations of transplantation experiments that focus on only white blood cell reconstitution (Schaefer *et al.*, 2001; Hamanaka *et al.*, 2013; Yamamoto *et al.*, 2013; Boyer *et al.*, 2019). Additionally, analysis of HSCs at the single cell level has identified an increased molecular Plt priming and functional Plt bias in the old HSC compartment (Sanjuan-Pla *et al.*, 2013; Grover *et al.*, 2016; Yamamoto *et al.*, 2018). These alterations raise the need to understand potential alterations in megakaryocyte differentiation in aged mice and humans.

The varying challenges to the blood system during aging may lead to alterations to multiple levels of differentiation, resulting in the dysregulation of homeostatic control maintained by committed progenitors. Previous studies have noted that the bone marrow (BM) frequencies of committed lymphoid and myeloid progenitors are altered with age (Morrison *et al.*, 1996; De Haan and Van Zant, 1999; Miller and Allman, 2003; Rossi *et al.*, 2005), but the functional properties and roles of hematopoietic progenitor cells during aging remains an enigma. Here, we investigated young and old HSCs and the contribution of both intrinsic changes and the environment on their differentiation. In our analysis, we included quantitative measurements of their erythroid and Plt production, as well as assessment of progenitor cells at steady-state and upon heterochronic HSC transplantation. These findings prompted us to further pursue age-associated changes in the Plt lineage, leading to unexpected revelations on the effect of aging on megakaryopoiesis.

## RESULTS

### Both the Frequencies and Numbers of Hematopoietic Stem and Progenitor Cell Populations are Altered During Aging

Aging has been shown to alter the frequencies of HSCs in the BM (Morrison *et al.*, 1996; De Haan and Van Zant, 1999; Sudo *et al.*, 2000; Rossi *et al.*, 2005). To evaluate if similar patterns were observed in the total cell numbers and of more refined erythromyeloid populations (Pronk *et al.*, 2007), we compared the absolute cell numbers of HSPC populations between young and old mice. Consistent with previous reports, the frequencies of the cKit+Lineage-Sca1+ (KLS) population increased dramatically with age. (**Figure 1A-B**). This increase was also observed when presented as total cell numbers (**Figure 1C**). When we further fractionated the KLS compartment, we found that this KLS abundance in the old BM was due to a dramatic enrichment of old HSCs. By contrast, the frequency and number of MPPs remained consistent throughout life. The Myeloid Progenitor (MyPro) compartment increased in total cell numbers, but not frequency. Classically defined CMPs and GMPs (Akashi *et al.*, 2000) were found at similar frequencies in young and old BM, while MEP frequencies were reduced with age (**Figure 1B**). Due to the overall increased cell numbers with age, however, absolute quantification revealed an increase in CMP and GMP numbers, but no change in MEPs, in old BM.

**Figure 1:**
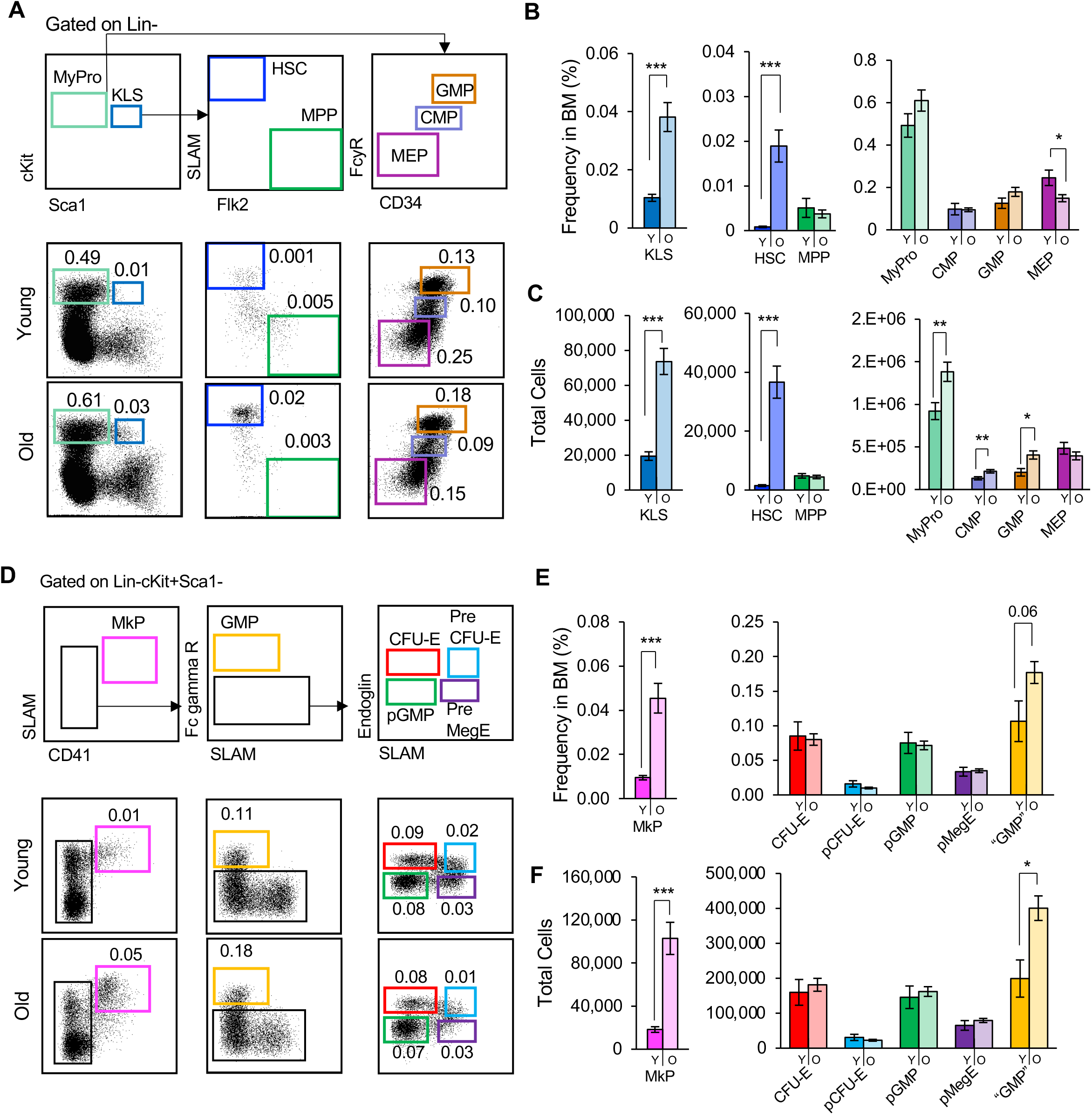
Hematopoietic Stem Cells and Megakaryocyte Progenitor Cells Expand During Aging. (A) Aging alters the composition of HSCs and classical myeloid progenitors. Gating and representative flow cytometry data of phenotypic HSCs, MPPs, and classical myeloid progenitor populations (phenotypic KLS, CMP, GMP, and MEP) in young and old bone marrow. (B-C) Increased frequencies and total cell numbers of HSCs of old mice compared to young mice. MPP cell percentages and total cell numbers remained the same in young and old mice. Frequencies of CMPs and GMPs remained unchanged while MEPs decreased in frequencies (B), but not numbers (C). The total cell numbers of MyPros, CMPs, and GMPs were significantly increased (C). (D) Aging alters the composition of erythromyeloid progenitors. Gating and representative flow cytometry data of MkPs and additional erythromyeloid progenitor cells (CFU-E, pCFU-E, pGMP, pMegE, “GMP”) in young and old bone marrow. (E-F) Frequency and total cell analysis revealed significantly higher levels of MkPs in the bone marrow of old mice compared to young mice. There was also a slight increase in the frequency and total cell numbers of “GMPs”. Y, Young; O, Old; HSC, Hematopoietic Stem Cells; MPP, Multipotent Progenitors; KLS, cKit+Lineage-Sca1+ cells; MyPro, Myeloid Progenitors; CMP, Common Myeloid Progenitors; GMP, Granulocyte/Macrophage Progenitor; MEP, Megakaryocyte-erythroid Progenitors; MkP, Megakaryocyte Progenitors. Additional abbreviations: Young n = 7, Old = 27 in 3 independent experiments. Data are shown as mean ± SEM. P values were determined using unpaired two-tailed t-test. *p<0.05, **p < 0.005, ***p < 0.0001.

We also investigated erythromyeloid populations more recently defined by alternative markers within the MyPro compartment (**Figure 1D**) (Pronk *et al.*, 2007). Few age-related alterations in frequencies and numbers were observed: CFU-E, pCFU-E, pGMP and pMegE frequencies and numbers were not significantly altered (**Figure 1E-F**). Alternatively defined “GMPs” trended towards an increase in frequency and were significantly increased in absolute numbers (**Figure 1E-F**), consistent with the highly overlapping classical GMPs (**Figure 1B-C**). Intriguingly, we observed a dramatic increase in both the frequency and total cell numbers of phenotypically defined MkPs in old mice **(Figure 1E-F).** These data revealed that the composition of HSPC populations selectively changes during aging. Of note was the dramatic expansion of old HSCs and old MkPs, raising the need to explore their role in megakaryopoiesis during the aging process.

### Old HSCs Exhibit Impaired Reconstitution Potential Compared to Young HSCs

To directly interrogate the differences in reconstitution potential between young and old HSCs, we purified HSCs from fluorescent young or old mice and transplanted them into young recipients (**Figure 2A)**. These experiments are analogous to those performed previously by others (Rossi *et al.*, 2005; Ergen *et al.*, 2012), except we expanded aforementioned investigations by also including direct detection of donor-derived Plts and circulating erythroid cells, in addition to the more commonly assayed granulocytes/myelomonocytes (GMs), and B and T cells.

**Figure 2:**
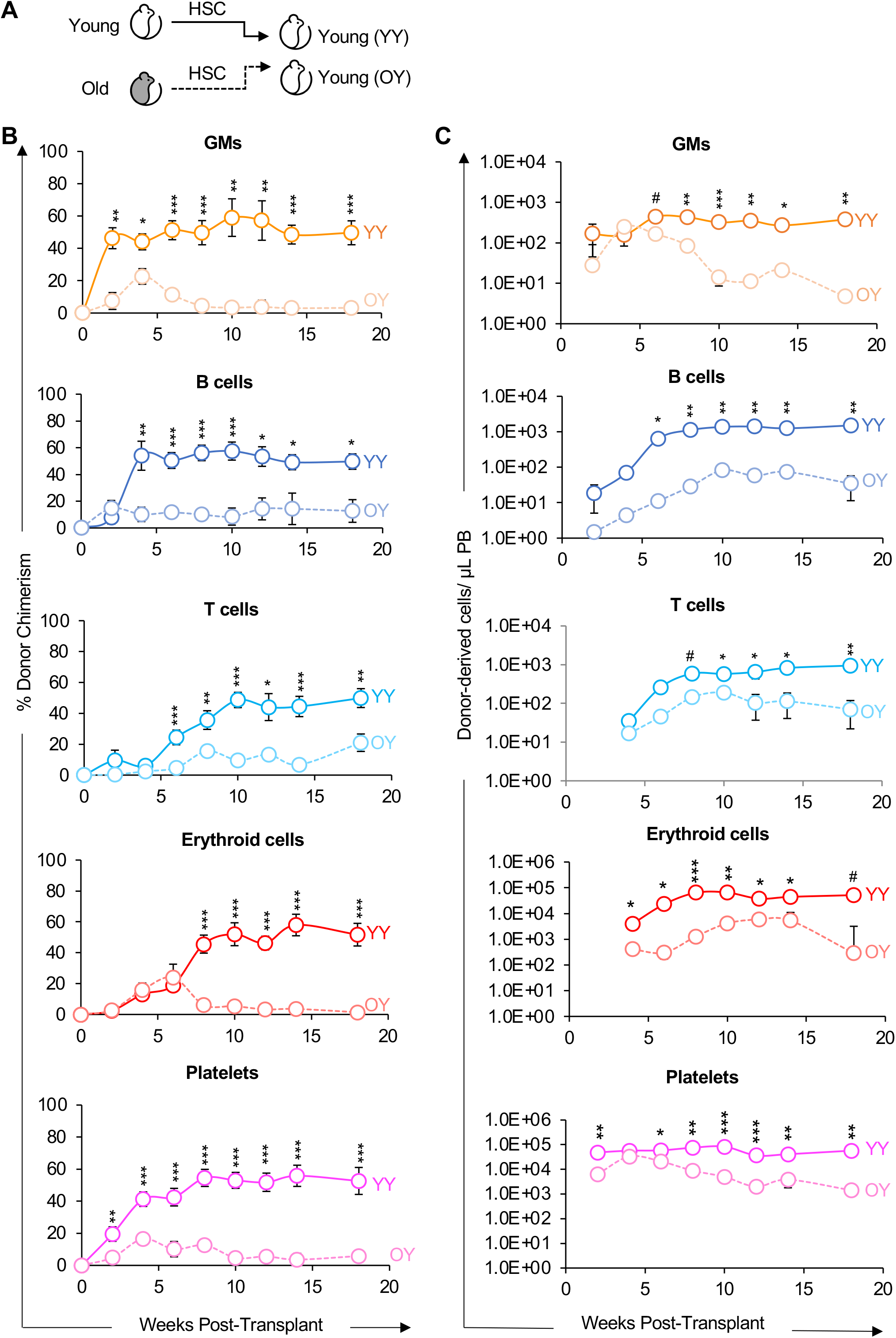
Old HSCs exhibited impaired reconstitution potential in young recipients. (A) Schematic of transplantation of HSCs from young or old mice into young recipient mice. (B) Old HSCs contributed to donor-host-chimerism with lower efficiency compared to young HSCs. Analysis of donor-derived mature cells in peripheral blood of recipients presented as percent donor chimerism. (C) Old HSCs produced fewer mature cells compared to young HSCs. Reconstitution data from (B) replotted as absolute numbers of donor-derived GMs, B cells, T cells, erythroid cells, and platelets per microliter of peripheral blood. GM, granulocytes/myelomonocytes. YY, Young HSCs transplanted into young recipients; OY, Old HSCs transplanted into young recipients. Data are representative means ± SEM from 9 recipient mice of young HSCs (4 independent experiments) and 22 recipients of old HSCs (5 independent experiments). P values were determined using unpaired two-tailed t-test. # p < 0.06, *p < 0.05, **p < 0.005, ***p < 0.0005.

First, also as commonly done, we determined the reconstitution kinetics of HSCs by measuring donor chimerism in the peripheral blood (PB) for the duration of 18 weeks post-transplantation (**Figure 2B**). Consistent with previous reports, analysis of lineage potential between young and old HSCs revealed that old HSCs have a diminished capacity to reconstitute GM, B, and T cells. Similarly, erythroid and Plt chimerism were also lower from old versus young HSCs (**Figure 2B**).

Second, we removed the host cell variable by quantifying the absolute numbers of donor-derived mature cells in the PB of recipient mice (Boyer *et al.*, 2019; Cool *et al.*, 2020; Rajendiran *et al.*, 2020a; Rajendiran *et al.*, 2020b) to more quantitatively assess the differences in regenerative activity between young and old HSCs. Old HSCs generated fewer erythroid, GM, B, T, and Plt cells compared to young HSCs (**Figure 2C**); this observation is consistent with host cell production presented as donor-derived chimerism (**Figure 2B**). Our data demonstrated a pan-hematopoietic reconstitution deficit of HSCs during aging, which is consistent with previous reports in white blood cell reconstitution. Taken together, these results also support the notion that key changes intrinsically maintained by HSCs drive their feeble performance upon aging.

### The Old Bone Marrow Environment Impairs the Reconstitution Potential of HSCs

We next sought to determine how cues in the aging environment influences HSC reconstitution capacity. Because young HSCs outperformed old HSCs in young hosts (**Figure 2**), we reasoned that young HSCs would engraft with higher efficiency in old compared to young recipients. To test this directly, we purified HSCs from young mice and transplanted them into young or old recipient mice (**Figure 3A**). We then analyzed mature cell production of transplanted HSCs to quantitatively assess the role of host age on expansion capacity and lineage potential. Young HSCs were capable of long-term, multilineage reconstitution (LTMR) in both young and old hosts. However, in contrast to the young hosts, the old hosts allowed for significantly lower reconstitution chimerism of HSC-derived GM, B, T, erythroid, and Plt cells (**Figure 3B**). Overall, the differences were more significant at earlier timepoints (~4-14 weeks) than at later timepoints.

**Figure 3:**
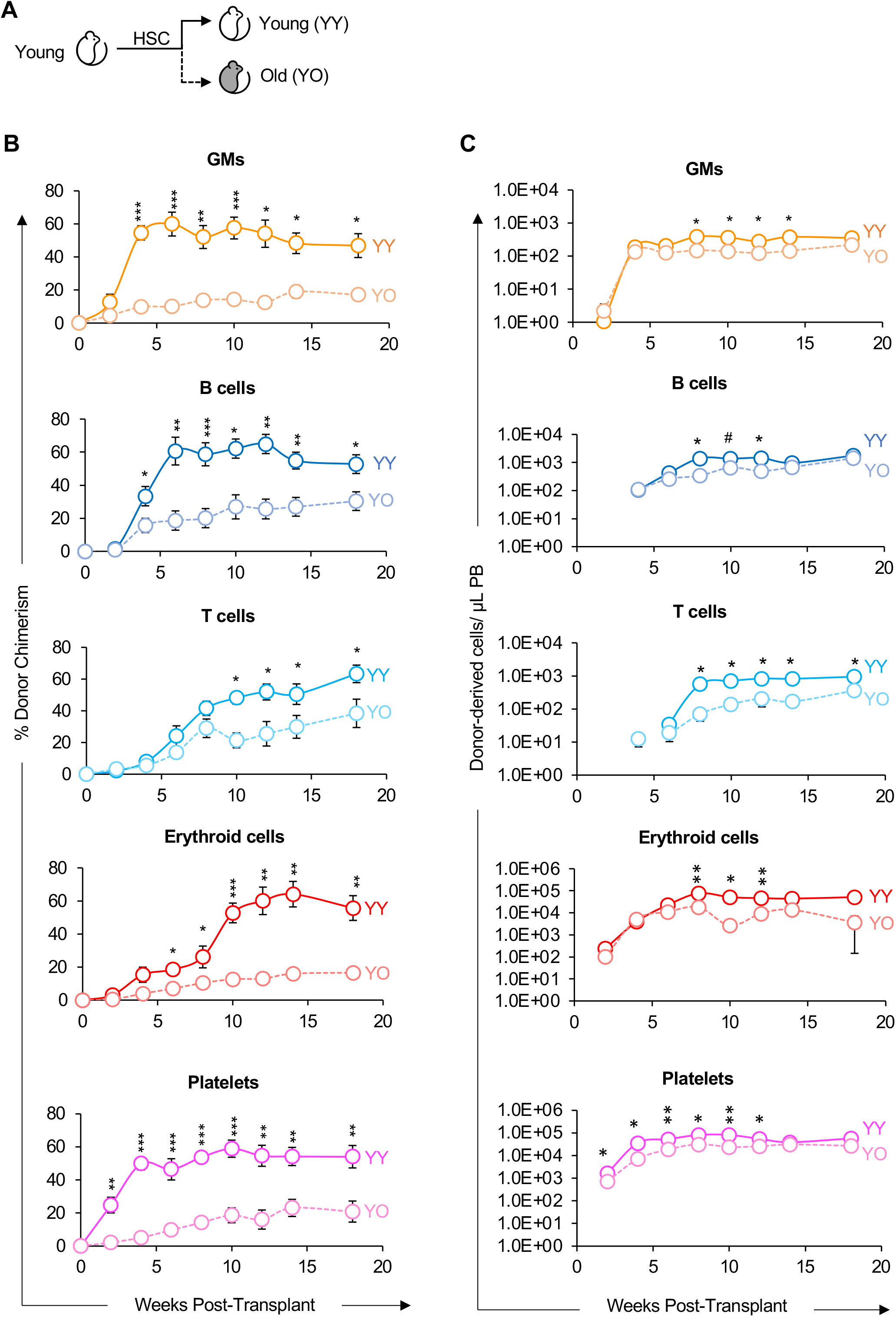
Young HSCs exhibited lower reconstitution potential in old compared to young recipients. (A) Schematic of transplantation of HSCs from young mice into separate cohorts of young or old mice. (B) Old recipients displayed lower HSCs donor-host-chimerism compared to young recipients. Analysis of donor-derived mature cells (GMs, B cells, T cells, erythroid cells, and platelets) in peripheral blood of recipients presented as percent donor chimerism. (C) Donor HSCs produced fewer total mature cells in old recipients compared to in young recipients. Reconstitution data from (B) replotted as absolute number of donor-derived GMs, B cells, T cells, erythroid cells, and platelets per microliter of peripheral blood. YY, Young HSCs transplanted into young recipients; YO, Young HSCs transplanted into old recipients. Data are representative means ± SEM from 9 young recipient mice (4 independent experiments) and 10 old recipient mice (3 independent experiments). P values were determined using unpaired two-tailed t-test. *p < 0.05, **p < 0.005, ***p < 0.0005.

In the same cohorts, we also determined the absolute number of donor-derived mature cells in the PB after transplantation. The donor HSCs in the young host produced greater numbers of donor-derived cells per microliter of PB compared to the old host (**Figure 3C**). Though this pattern was overall consistent with the patterns observed by donor-chimerism, the differences in total cell production was attenuated and, with the exception of T cells, was not sustained over time. This analysis revealed that the old host reduced donor-chimerism by young HSCs. However, when we removed the host variable by measuring absolute mature cell production, our analysis revealed a less different reconstitution capacity between young HSCs placed in the young environment and the young HSCs in the old environment.

Because the isochronic Y-into-Y transplants led to higher chimerism than the two heterochronic (O-into-Y or Y-into-O) transplants (**Figures 2 and 3**), we hypothesized that an old environment would provide a more supportive environment for old HSCs. We therefore compared the reconstitution potential of young and old HSCs in an old host (**Figure 4A**) and assayed for donor-chimerism (**Figure 4B**) and absolute number of donor-derived cells (**Figure 4C**). Surprisingly, transplantation of young HSCs led to higher reconstitution of old recipients compared to transplantation of old HSCs. We also examined the impact of reconstitution by old HSCs in a young or old environment; our analysis revealed that old HSCs exhibited poor reconstitution capacity in the old host compared to the young host (**Figure S1**). Therefore, while the aged BM environment impaired the reconstitution potential of transplanted young HSCs (**Figure 3**), HSCs from old mice performed even worse than young HSCs in an aged environment (**Figure 4, Figure S1-2A-A”‘**). In combination with the diminished reconstitution potential of old HSCs (**Figure 2**), these results indicate that aging also leads to reduction in HSC-intrinsic performance and in the ability of the BM environment to support transplanted HSCs (**Figure 3, 4, S1,S2**).

**Figure 4:**
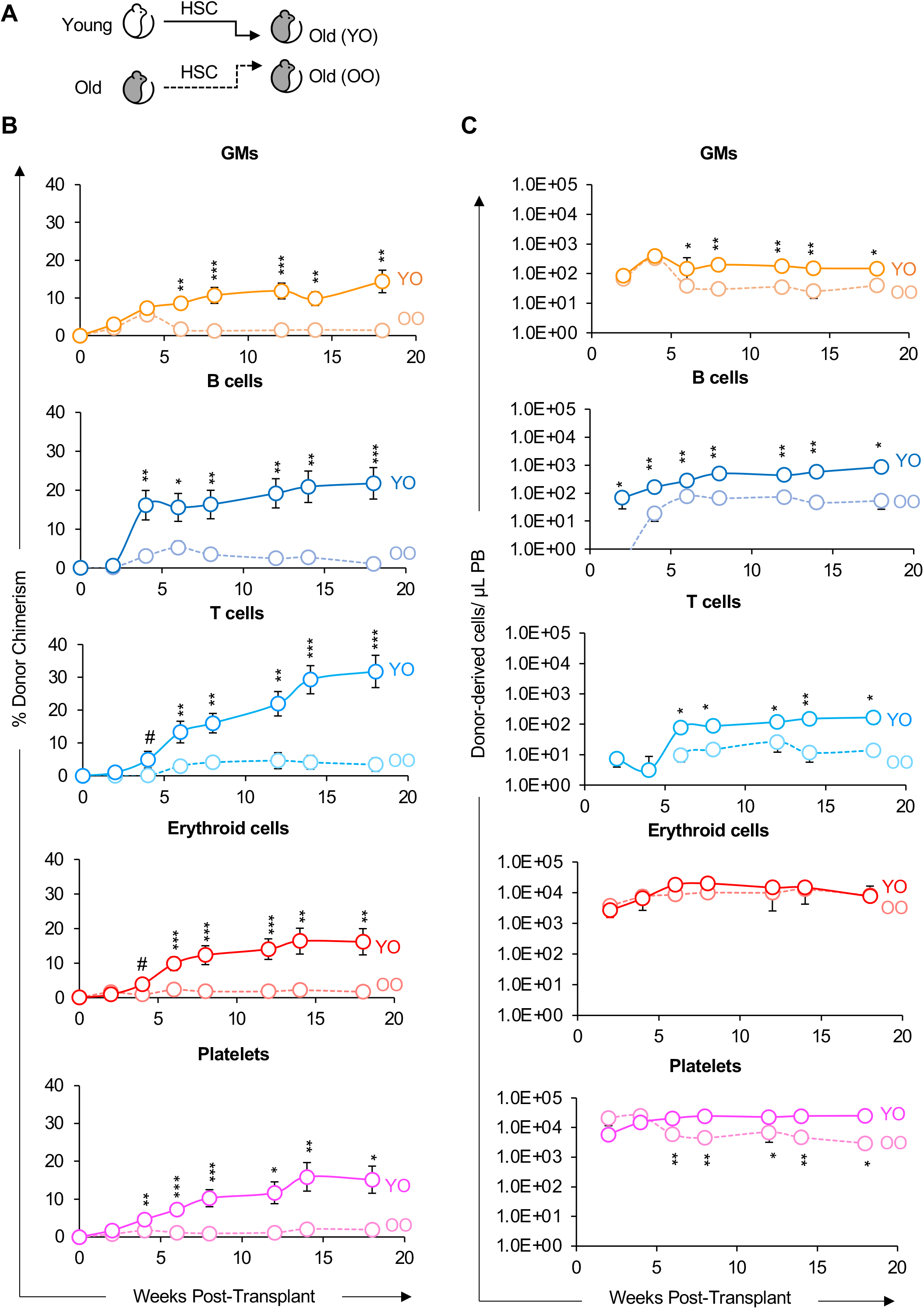
Old HSCs exhibited impaired reconstitution potential compared to young HSCs in old recipients. (A) Schematic of transplantation of HSCs from young or old mice into separate cohorts of old mice. (B) Old HSCs displayed lower donor-host-chimerism compared to young HSCs. Analysis of donor-derived mature cells (GMs, B cells, T cells, erythroid cells, and platelets) in peripheral blood of recipients presented as percent donor chimerism. (C) Old HSCs produced fewer total mature cells in old recipients compared to in young HSCs. Reconstitution data from (B) replotted as absolute number of donor-derived GMs, B cells, T cells, erythroid cells, and platelets per microliter of peripheral blood. YO, Young HSCs transplanted into old recipients; OO, Old HSCs transplanted into old recipients. Data are representative means ± SEM from 11 recipient mice of young HSCs (4 independent experiments) and 8 recipients of old HSCs (3 independent experiments). P values were determined using unpaired two-tailed t-test. *p < 0.05, **p < 0.005, ***p < 0.0005.

### Both old HSCs and the old niche impairs reconstitution of the stem and progenitor cell compartment

The reported Plt bias of old HSCs (Sanjuan-Pla *et al.*, 2013; Yamamoto *et al.*, 2013, 2018; Haas *et al.*, 2015; Grover *et al.*, 2016) had the potential to translate into maintained, or even increased, Plt reconstitution by old HSCs, possibly at the expense of the other lineages. However, Plt reconstitution by old HSCs in either young (**Figure 2**) or old (**Figure 4**) hosts, and by young HSCs in old hosts (**Figure 3**) was as impaired as reconstitution of erythroid cells and WBCs. To determine whether the observed reconstitution of the PB is reflected in the BM of the recipients, we also determined the capacity of young and old HSCs to regenerate stem and progenitor cells. In the cohorts analyzed for PB reconstitution in Figures 2 and 3, we determined donor chimerism in the BM stem and progenitor cell compartment >18 weeks post-transplantation (**Figure 5A**). All expected populations were detected, supporting the long-term multilineage reconstitution (LTMR) capability of HSCs. Quantitatively, young recipients contained similar total BM cell numbers (**Figure 5A’**) and, in comparison to young HSCs, old HSCs exhibited a much lower capacity to reconstitute classical HSPCs and erythromyeloid progenitor populations, with notably lower contribution to MkP reconstitution (**Figure 5A”**). Likewise, quantifying the total cell number of donor-derived HSPCs revealed a similar pattern (**Figure 5A”‘**). When determining the role of the aging recipient environment on HSC reconstitution of BM populations (**Figure 5B, S2B-B’’’**), we also found that despite having similar total BM cell numbers (**Figure 5B’ and S2B’**), there were lower frequencies and total numbers of donor-derived classical myeloid progenitor cells and erythromyeloid progenitor populations in the old hosts compared to the young hosts (**Figure 5B’-B’’**). Similar observations were made for the old HSCs in old recipients (**Figure S2A-A’’’**). Taken together, our data suggest that aging alters both HSC-intrinsic and recipient properties, and that these age-related changes affect the entire hematopoietic hierarchy and all lineages, including erythro- and megakaryopoiesis.

**Figure 5:**
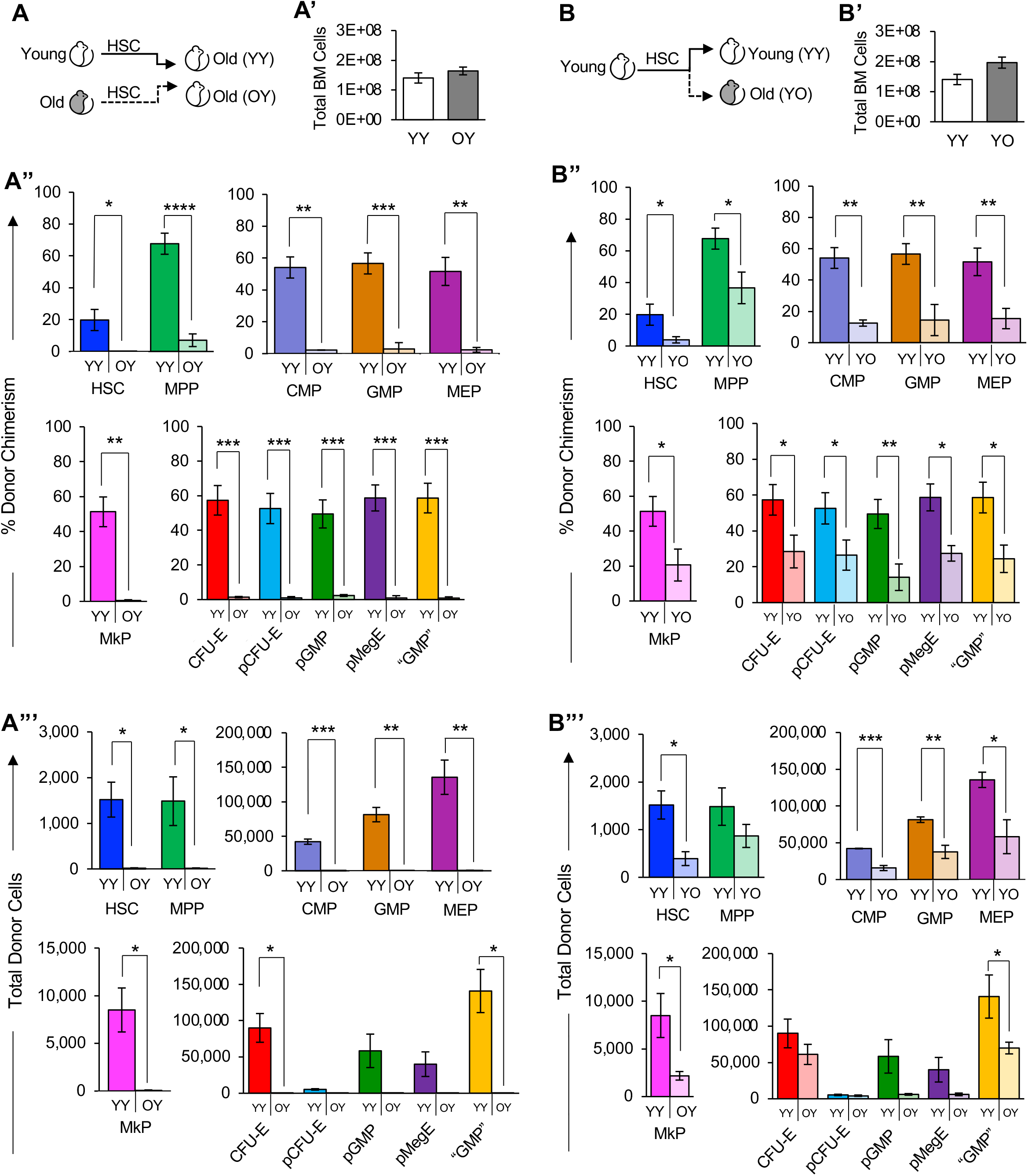
Both old HSCs and the old niche impaired reconstitution of the stem and progenitor cell compartment. (A) Schematic of HSC transplantation from young or old mice into young mice. (A’) The total number of bone marrow cells were similar in young recipient mice >16 wks after transplantation of either young or old HSCs. (A’’) Old HSCs contributed to significantly lower donor-to-host chimerism of bone marrow stem and progenitor cells in young recipient mice compared to young HSCs. Percent donor chimerism of HSPCs and erythromyeloid progenitors in the recipient bone marrow of young or old HSCs > 16 weeks post-transplant. (A’’’) Old HSCs generated fewer total stem and progenitor cells than young HSCs in young recipient mice. Quantification of total donor-derived classical HSPCs and erythromyeloid progenitors in the bone marrow of recipients > 16 weeks post-transplant. (B) Schematic of HSC transplantation from young mice into separate cohorts of young or old mice. (B’) The total number of bone marrow cells were similar in young and old recipient mice >18 wks after transplantation of young HSCs. (B’’) Old recipients displayed lower donor-to-host HSPC chimerism than young recipient mice transplanted with young HSCs. BM analysis of young or old mice transplanted with Young HSCs. HSPCs were analyzed > 18 weeks post-transplant and presented as donor chimerism. (B’’’) Transplanted young HSCs produced fewer total HSPCs in the bone marrow of old compared to young recipient mice. Quantification of total donor-derived HSPC numbers in the bone marrow of recipients > 18 weeks post-transplant. YY, Young HSCs transplanted into young recipients; OY, Old HSCs transplanted into young recipients; YO, Young HSCs transplanted into old recipients. Data are representative means ± SEM. PB analysis of the same recipient mice are presented in Figure 2 (panels A’-A’’’) and Figure 3 (panels B’-B’’’). P values were determined using unpaired two-tailed t-test. *p≤0.05, **p<0.005, ***p<0.0005, ****p<0.0001.

### Old MkPs were Functionally Enhanced Compared to Young MkPs

The most profound and consistent change in old mice was alterations in megakaryopoiesis, denoted by a dramatic increase in MkPs (**Figure 1**). However, this age-dependent increase in MkPs was not accompanied by an age-dependent functional bias of HSCs towards the megakaryocyte lineage: transplanted old HSCs were as deficient in MkP (**Figures 5A’’-B’’, A’’’-B’’’, S2A’’-B’’, S2A’’’-B’’’**) and Plt (**Figures 2B-C, 4B-C, S1B-C**) reconstitution as they were in generating other lineages. Thus, we hypothesized that the dramatic increase in the frequency and total numbers of old MkPs (**Figure 1D-F)** may be manifested by MkP-intrinsic changes, possibly in addition to increased contribution from the HSC compartment.

To begin to test this hypothesis, we determined functional differences between young and old MkPs *in vitro.* Interestingly, evaluation of MkP proliferative capacity by short-term culture assays revealed a significant, 2.5-fold increase in expansion of old MkPs compared to young MkPs (**Figures 6A-B**). These experiments showed that old MkPs displayed a surprising *increased* functional capacity compared to young MkPs *in vitro.*

**Figure 6:**
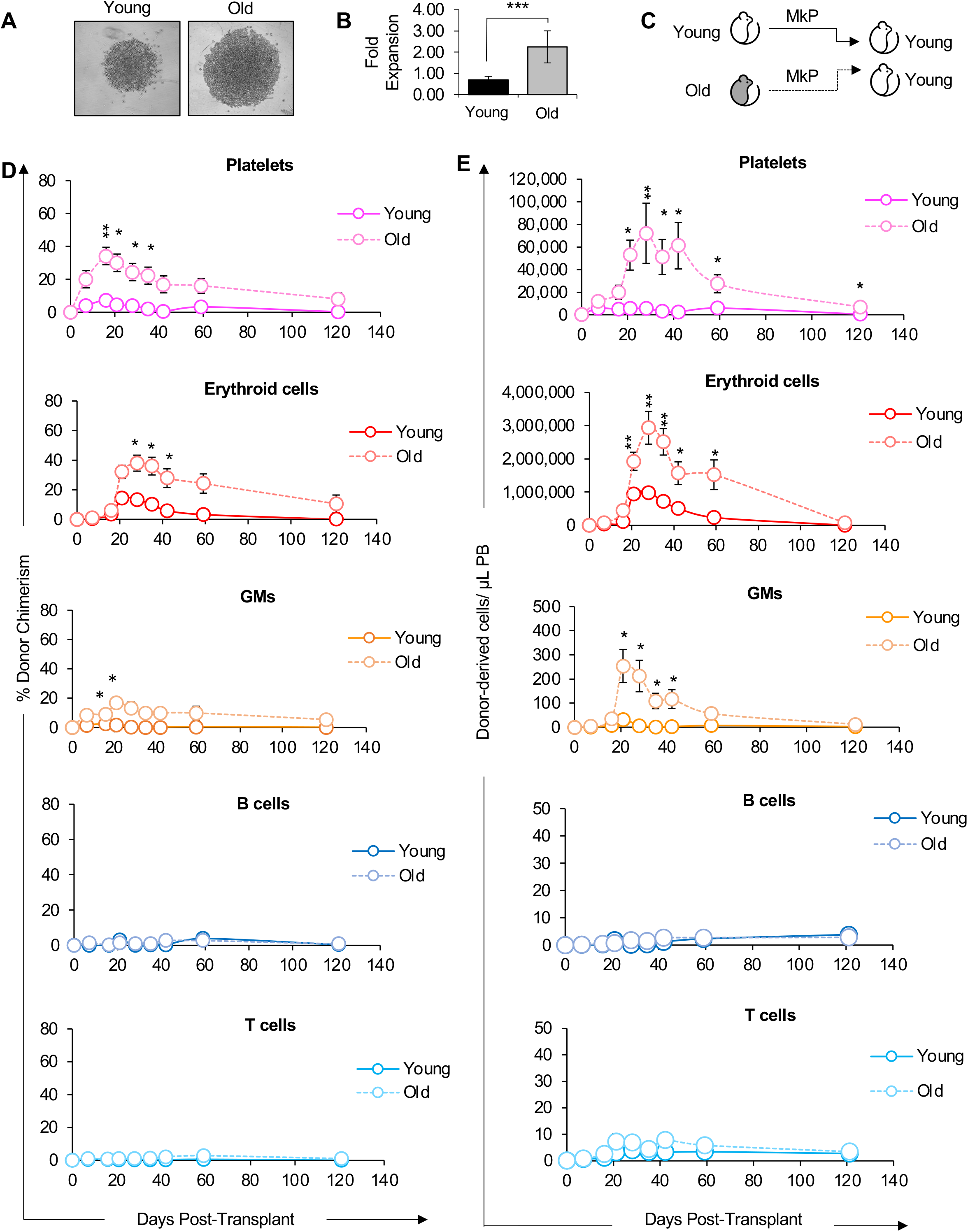
Old MkPs had enhanced reconstitution potential compared to young MkPs. (A-B) Old MkPs have enhanced proliferative capacity compared to young MkPs *in vitro.* (A) Representative images of Y and O MkPs grown *in vitro.* To evaluate proliferative capacity, 2,000 MkPs were FACS sorted from young and old mice and grown in liquid culture. 3 days after seeding, MkPs were microscopically evaluated for expansion efficiency. (B) MkP expansion *in vitro.* Flow cytometry was used to quantify the total fold expansion of cultured young and old MkPs from (A). n=3 from 3 independent experiments. (C) Schematic of transplantation of young or old MkP into young mice. (D) Old MkPs demonstrated greater short-term contribution to platelet donor-to-host chimerism in the recipient mice compared to young MkPs. Old MkPs also contributed to short-term erythroid cells and GM donor-to-host chimerism, but not to B or T cells. Analysis of donor-derived mature cells (platelets, erythroid cells, GMs, B cells, T cells) in peripheral blood of recipients presented as percent donor chimerism. (E) Old MkPs generated higher numbers of platelets compared to young MkPs in recipient mice. Old MkPs also produced more erythroid cells and GMs compared to young MkPs. Reconstitution data replotted as absolute numbers of donor-derived platelets, erythroid cells, GMs, B cells, and T cells per microliter of peripheral blood. Data are representative means ± SEM. Data from transplantation experiments are from 5 recipient mice of young MkPs and 11 recipient mice of old MkPs from at least three independent experiments. P values were determined by unpaired two-tailed student’s t-test *p<0.05, **p<0.005, *** p≤0.001.

### Old MkPs Possessed Greater Platelet Reconstitution Potential Compared to Young MkPs

The notable increase in expansion capacity of old MkPs *in vitro* lead us to predict potential functional differences *in vivo.* To determine the reconstitution potential of old MkPs *in vivo*, we transplanted young or old MkPs into young recipient mice (**Figure 6C**). We evaluated MkP multilineage reconstitution in the host by measuring their contribution to mature cells (Plt, erythroid cells, GMs, B, T) in the PB for 16-weeks. Given that young MkPs have been shown to be strongly committed to the Plt lineage in *in vitro* assays (Nakorn *et al.*, 2002; Pronk *et al.*, 2007), we predicted that MkPs would primarily produce Plts. Indeed, young MkPs contributed to nominal donor-chimerism levels of GM, B, and T cells, with a more robust, contribution to Plts, as well as, unexpectedly, a short burst of erythroid cells (**Figure 6D**). Surprisingly, relative to young MkPs, old MkPs exhibited a remarkable capacity to engraft and contribute to Plts in the recipient mice (**Figure 6D**). While young MkPs repopulated a more modest 7.4% of Plts, the same number of old MkPs repopulated on average 34% of Plts in the recipient mice. Interestingly, old MkPs also contributed to significantly higher erythroid cell and GM chimerism in the recipient mice relative to young MkPs. It is also notable that unlike the long-term reconstitution capacity of HSCs (**Figures 2-5**), the Plt, erythroid cell, and GM production from Old MkPs was robust only in the first few weeks post-transplant and then declined over time.

These differences between young and old MkPs were reinforced by quantification of the absolute cell numbers from the same experiments (**Figure 6E**). The absolute quantification revealed that old MkPs engrafted and produced Plts, as well as erythroid cells and GMs, at a remarkably high efficiency compared to young MkPs. Taken together, our *in vitro* and *in vivo* results indicated that MkPs gain both expansion and reconstitution capacity during aging.

### RNA-sequencing revealed different molecular patterns between Young and Old MkPs

We were surprised by the remarkable difference in expansion and reconstitution capacity between young and old MkPs. To test how aging affects the molecular profile of MkPs, we evaluated the molecular regulation of old MkPs by performing RNA sequencing (RNAseq) analysis of young and old MkPs. We found that young and old MkPs had very similar gene expression profiles as expected of two MkP populations, sharing over 15,000 transcripts. Importantly, we also identified 520 differentially expressed genes between young and old MkPs, 237 of which were downregulated and 283 were upregulated in old MkPs (**Figure 7A**). Hierarchical clustering of seven independent samples showed that independent biological replicates associated based on age, as expected (**Figure 7B**), with the top 50 differentially expressed genes shown as a heatmap (**Figure 7B and Table S1**).

**Figure 7:**
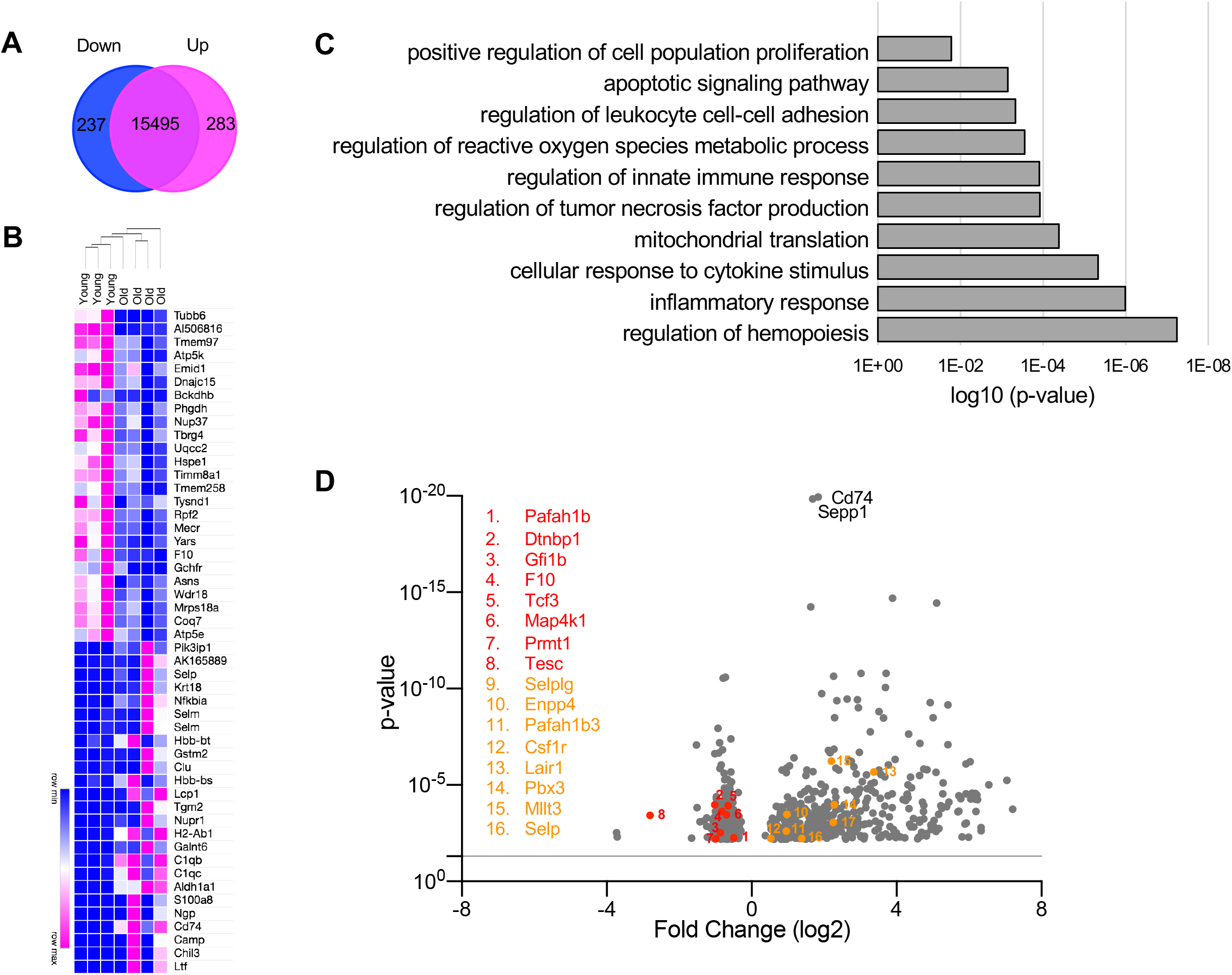
Aging is associated with a change in MkP genome-wide expression signatures. (A) RNAseq analysis revealed transcriptional changes accompanying aging of the MkP compartment. 237 genes were downregulated and 283 genes were upregulated in Old MkPs. A venn diagram showing the numbers of downregulated (blue) and upregulated (pink) DEGs in old versus young MkPs. (B) Hierarchical clustering analysis confirmed the distinct gene expression characteristics of the young and old MkPs. Heatmap showing the scaled expression of the top 50 differentially expressed genes (DEGs) in young and old MkP clusters. Pink and blue indicate comparatively high and low expression, respectively. (C) The expression patterns associated with aging MkPs revealed changes to important biological functions. Major Gene Ontology (GO) biological process terms in which DEGs enriched for. (D) Identifying specific DEGs highlight known and novel regulators of aging megakaryopoiesis. A volcano plot of DEGs between young and old MkPs. Genes described in text are highlighted.

To further evaluate the expression signatures of young and old MkPs, we performed Gene Ontology (GO) analysis. This analysis indicated that differentially expressed genes between young and old MkPs were highly enriched in categories related to proliferation (**Figure 7C**), inferring intrinsic regulation of the observed enhanced expansion capacity of old MkPs (**Figure 6A-E**). Other GO terms associated with the differentially expressed genes included cell adhesion, inflammation, and mitochondrial regulation (**Figure 7C**). The latter is consistent with a recent publication of mitochondrial activity playing a role in age-related Plt function (Davizon-Castillo *et al.*, 2019). Genes under each GO term are presented in **Table S2**. At the gene level, we observed both novel and well-known regulators of hematopoiesis and hematological processes (**Figure 7B,D**). For example, we identified regulators of lineage specification, including Csf1 and Tcf3 (also known as E2A) (Semerad *et al.*, 2009; Stanley and Chitu, 2014). Also, previously described regulators of megakaryocyte differentiation were differentially expressed. These genes include Prmt1, Selp, and Tesc, a regulator of E26 transformation-specific (ETS) transcription factors (Levay and Slepak, 2007; Jin *et al.*, 2018; Banu, Avraham and Karsenty, 2020). We also observed differential expression of multiple genes associated with Plt regulation, production, and function: Coagulation Factor X, Dtnbp1, Gfi1b, Enpp4, Pafah1b3, Pafah1b1, Mmrn1, and Selplg. Mutations in Gfi1b and Dtnbp1 are associated with various Plt disorders and bleeding disorders (Ghiani and Dell’Angelica, 2011; Songdej and Rao, 2017). Interestingly, our analysis also revealed a small subset of upregulation genes previously implicated in acute myeloid leukemia (AML), including Pbx3, Lair1, and Mllt3 (Marschalek, 2010; Li *et al.*, 2013; Kang *et al.*, 2015), and is possibly linked to increased risk of AML in the elderly (Almeida and Ramos, 2016).

The results of the RNAseq analysis corroborates the behaviors observed in our functional analyses of MkPs. Together, these data illuminate the fundamental difference between youthful and aging megakaryopoiesis. Importantly, our RNAseq data revealed well-known genes implicated in the molecular mechanisms of megakaryopoiesis during aging, as well as a number of novel genes that can be pursued to understand, and possibly mitigate, the age-related changes that occur in megakaryopoiesis.

### DISCUSSION

#### Aging alters the reconstitution potential of HSCs

As previously reported by others, our characterization of the old BM populations revealed an expansion of HSCs (**Figure 1**). This observation raised the need understand how HSC differentiation is modulated by the balance of both intrinsic changes to HSCs and extrinsic influence on HSCs during aging. Young HSCs have been suggested to have superior engraftment and reconstitution capacity compared to old HSCs (Morrison *et al.*, 1996; Sudo *et al.*, 2000; Rossi *et al.*, 2005; Dykstra *et al.*, 2011). Indeed, our experiments transplanting young or old HSCs indicated that old HSCs exhibited a diminished reconstitution capacity (**Figure 2 and 5**). To address whether the reduced reconstitution activity of HSCs during aging is influenced by the aged environment, we transplanted young HSCs into young or old hosts (**Figure 3 and 5**). Our data revealed that the old environment was less supportive of reconstitution by transplanted HSCs compared to the young environment. Therefore, transplanted young HSCs did not have an obvious competitive advantage over the resident old HSCs in the aged host. Notably, transplanted old HSCs had even lower capacity to reconstitute the old host compared to young HSCs (**Figure 4, S2**), demonstrating that the poor performance of old HSCs in young recipients (**Figure 2 and 5**) was not due to the age-mismatch (heterochronic) between donor and host. Because both young and old HSCs perform worse in old compared to young recipients, it is clear that age-related changes in the host impact donor HSC performance. This could be related to the reported increased proliferation of old HSCs (Kirschner *et al.*, 2017), age-related changes to the distal and local microenvironment, or both. Our collective iso- and heterochronic transplantation data show that both HSC-intrinsic and extrinsic mechanisms during aging affect reconstitution efficiency (SanMiguel *et al.*, 2020). Interestingly, we did not detect a selective retention of MkP or Plt reconstitution capacity of transplanted old HSCs, despite the increased frequencies and numbers of HSCs and MkPs in unmanipulated aged mice (**Figure 1**). It remains possible that HSCs contribute to the greater production of MkPs *in situ* (Carrelha *et al.*, 2018). Together with the observed expansion of HSCs *in situ* (**Figure 1**), our transplantation results suggest that the impaired reconstitution performance of aged HSCs reflects reduced engraftment capacity rather than functional decline *in situ.* A better understanding of how young and old BM cells respond to niche-clearing methods could help to develop optimal systems for investigating reconstitution differences between young and old HSCs, and between young and old recipients.

#### Surprising Reconstitution and Differentiation Potential of Old MkPs

Our data provided evidence about the changes to the homeostatic control of blood cell production at the MkP level of hematopoietic differentiation during aging. Given the reconstitution deficit displayed by old HSCs (**Figures 2, 4, 5A-A’’’**), one might have expected that old MkPs would also display functional deficiencies. We were surprised to find from our *in vitro* experiments that old MkPs displayed greater proliferative potential (**Figure 6A-B).** Strikingly, we also observed from transplanting young and old MkPs that old MkPs harbored a remarkable capacity to engraft, expand, and reconstitute Plts (**Figure 6C-E**). This age-associated elevation of MkP potential is reminiscent of the higher prevalence of thrombosis and other cardiovascular problems in the elderly. We hypothesize that age-related alterations made to MkP populations contribute to dysregulation and increased risk for age-related morbidities (Davizon-Castillo *et al.*, 2019).

Our transplantation studies of old and young MkPs also revealed somewhat surprising evidence about the differentiation programs of immature progenitors during aging (**Figure 6C-E**). Previous reports suggested that young MkPs are exclusively associated with megakaryocyte and Plt generation (Nakorn *et al.*, 2002; Pronk *et al.*, 2007). However, those conclusions were primarily based on transcription profiles and *in vitro* differentiation capacity, with no direct assessment of differentiation potential *in vivo.* We recently reported that classically (Akashi *et al.*, 2000) and alternatively (Pronk *et al.*, 2007) defined GMPs produce erythroid cells *in vivo.* This fits a model where erythropoiesis is a default HSC fate (Boyer *et al.*, 2019). Under that model, it is not surprising that our experiments revealed that both young and old MkPs harbor *in vivo* erythroid capacity. Surprisingly, old MkPs also demonstrated greater capacity for the GM lineage compared to young MkPs. It is possible that our observations may be a result of poorly defined cellular phenotypes in old mice, raising the question of how aging might affect the immunophenotype of MkPs. Given that several markers are shared between MkPs and HSCs (Zhu *et al.*, 2019), with a gain of CD41 expression as HSCs age (Gekas and Graf, 2013) the possibility that the MkP compartment in old BM contains significant numbers of HSCs was considered. However, old MkPs lacked B and T-cell generation and the enhanced production of Plts, erythroid cells, and GMs did not persist long-term, unlike that of multipotent and self-renewing HSCs. Additionally, we did not observe any age-related differences in expression of megakaryopoiesis master regulators such as Fli1, Gabpa, Mpl, Pf4, and Runx1 (De Sauvage *et al.*, 1994; Kaushansky *et al.*, 1994; Elagib *et al.*, 2003; Klimchenko *et al.*, 2009; Doré and Crispino, 2011; Bruns *et al.*, 2014; Zhu *et al.*, 2018) (**Figure 7**). A high similarity between these genes suggest high purity of the MkP populations. Together, these findings indicate that the high expansion and oligopotency of old MkPs by transient production of Plts, erythroid cells, and GMs cannot be explained by contaminating HSCs, but rather due to enhanced potential in the aged MkP compartment.

#### Aging is associated with changes in MkP genome-wide expression signatures

Our transcriptome data provide important clues about the intrinsic molecular regulation of megakaryopoiesis during aging (**Figure 7**). RNAseq analysis of young and old HSCs has provided a paradigm of molecular switches during ontogeny (Sanjuan-Pla *et al.*, 2013; Cabezas-Wallscheid *et al.*, 2014; Sun *et al.*, 2014; Kowalczyk *et al.*, 2015; Grover *et al.*, 2016). Similarly, our data showed that young and old MkPs have different gene expression programs, reflecting divergence in the molecular control of Plt differentiation during aging. For example, differential expression of genes involved in proliferation reinforced the differences in proliferative capacity between young and old MkPs. The expansion of MkPs with age also suggest that MkPs themselves may serve as a reservoir for mutations promoting clonal progression to cardiovascular diseases and cancer (Pardali *et al.*, 2020; Steensma and Ebert, 2020). In line with this model, we observed age-related increase in expression of genes involved in AML. Moreover, different transcriptional networks involving Plt function were observed between young and old MkPs, supporting a model in which MkPs propagate events towards age-related dysregulation of Plt biology. An important next step will be to identify key molecular regulators that cause or are a consequence of age-related changes to megakaryopoeisis.

In conclusion, our data revealed that profound perturbations in homeostatic control underlie megakaryopoiesis during the aging process. Given that HSCs gradually lose their regenerative potential during aging, it is reasonable to focus research on the HSC contribution to aging. There remains, however, a conceptual gap in the role of altered progenitor cells as potential perpetuators of aging. Our findings shed light on the mechanisms of aging within the Plt lineage that may dictate the etiology of Plt-related disorders, and other myeloid diseases that may be operating through committed progenitors rather than HSCs. Our study sets the stage for further exploration of mechanisms governing youthful and aging megakaryopoiesis and suggests that progenitor cell involvement should be considered when investigating normal and pathophysiological changes in the hematopoietic system during aging.

### MATERIALS AND METHODS

#### Mouse lines

All animals were housed and bred in the AAALAC accredited vivarium at UC Santa Cruz and maintained under approved IACUC guidelines. The following mice were utilized for these experiments: C57Bl6 (JAX, cat# 664), aged C57Bl6 (NIH-ROS), BoyJ (JAX, cat# 2014), Ubc-GFP (JAX, cat# 4353). Young adult mice were used between 8 – 16 weeks of age and old adult mice were 20+ months of age. Old recipient mice in transplantation experiments were 18+ of age. All mice were randomized based on sex.

#### Flow Cytometry

Bone marrow stem and progenitor cell populations and mature cell subsets were prepared and stained as previously described (Beaudin, Boyer and Forsberg, 2014; Smith-Berdan *et al.*, 2015, 2019; Ugarte *et al.*, 2015; Beaudin *et al.*, 2016; Leung *et al.*, 2019; Martin *et al.*, 2020). Briefly, the long bones (femur and tibia) from mice were isolated, crushed with a mortar and pestle, filtered through a 70μm nylon filter and pelleted by centrifugation to obtain a single cell suspension. APC-conjugated spherobeads (BD Bioscience) were added to the cell suspension prior to staining cells with fluorescently labeled antibodies to cell surface antigens. Cell labeling was performed on ice in 1X PBS with 5 mM EDTA and 2% serum. The stem and progenitor populations within the bone marrow were characterized as: HSC: Lin^-^/ckit^+^/Sca1^+^/Flk2^-^/Slam^+^, MPP: Lin^-^/ckit^+^/Sca1^+^/Flk2^+^/Slam^-^, MkP: Lin^-^/ckit^+^/Sca1^-^/CD41^+^/Slam^+^, CMP: Lin^-^/ckit^+^/Sca1^-^/FcγR^mid^/CD34^mid^, GMP: Lin^-^/ckit^+^/Sca1^-^/FcγR^hi9h^/CD34^hi^≡^h^, MEP: Lin7ckit7Sca17FcγR7CD34^-^, Pre-GM (Pronk): Lin^-^/cKit^+^/Sca1^-^/CD41^-^/ FcγR7CD1507Endoglin^-^, Pre-Meg (Pronk): Lin^-^/cKit^+^/Sca1^-^/CD41^-^/ FcγR7CD1507Endoglin^-^, Pre-CFU-E (Pronk): Lin7cKit7Sca17CD41^-^/ FcγR^-^ /CD150^+^/Endoglin^+^. Mature cells were characterized by: erythroid cells as circulating erythrocyte progenitors: (FSC^lo-mid^/Ter-119^+^/CD71^+^/Mac1^-^/Gr1^-^/B220^-^/CD3) or erythrocytes: (FSC^lo-mid^/Ter-119^+^/CD61^-^/Mac1^-^/Gr1^-^/B220^-^ /CD3^-^), platelets: (SSC7 Ter-119^-^/CD61^+^/Mac1^-^/Gr1^-^/B220^-^/CD3^-^), GM (Ter-119^-^/CD61^-^/Mac1^+^/Gr1^+^/B220^-^/CD3^-^), B-cell (TER-1197CD617Mac17Gr17B220^+^/CD3^-^), T-cell (Ter-119^-^/CD61^-^/Mac1^-^/Gr1^-^/B220^-^/CD3^+^). Cell suspensions were analyzed for specific cell populations using FACS Aria III (Becton Dickinson, San Jose, CA).

The lineage cocktail was comprised of CD3 (Biolegend cat #100306), CD4 (Biolegend cat #100423), CD5 (Biolegend cat #100612), CD8 (Biolegend cat #100723), Ter-119 (Biolegend cat #116215), Mac1 (Biolegend cat #101217), Gr1 (Biolegend cat #108417), and B220 (Biolegend cat #103225). Antibodies used in sorting were: cKit(Biolegend cat #105826), Sca1 (Biolegend cat #122520), CD150 (Biolegend cat #115914), FLK2 (ebiosciences cat #12-1351-83), CD34 (ebiosciences cat #13-0341-85), CD41 (Biolegend cat #133914), CD105 (Biolegend cat #120402). Antibodies used in peripheral blood were: CD71 (Biolegend cat # 113803), CD61 (Biolegend cat # 104314), CD3 (Biolegend cat #100306), TER-119 (Biolegend cat #116215), Mac1 (Biolegend cat #101217), Gr1 (Biolegend cat #108417), and B220 (Biolegend cat #103225).

#### HSC and MkP sorts by Flow Cytometry

HSC (Lin^-^/cKit^+^/Sca1^+^/Flk2^-^/Slam^+^) or MkP (Lin^-^/cKit^+^/Sca1^-^/CD41^+^/Slam^+^) from young or old mice were prospectively isolated using a FACS ARIA III (Becton Dickinson, San Jose, CA) as previously described [ref]. Cells were harvested from long bones as mentioned above and stained with CD117-microbeads (Miltenyi), then passed over a magnetic column to enrich for CD117^+^ stem and progenitor cells. HSC and MkP cell populations were double sorted on low pressure with a 100μm nozzle into PBS, 2% serum.

#### Transplantation Reconstitution assays

Reconstitution assays were performed by transplanting double-sorted HSCs (200 per recipient) from young or old Ubc-GFP^+^ whole bone marrow and transplanting into congenic C57Bl6 mice via retro-orbital intravenous transplant. We also transplanted double-sorted MkPs (22,000 per recipient) from C57Bl6 into Ubc-GFP^+^ hosts. Hosts were preconditioned with sub-lethal radiation (~750 rads) using a Faxitron CP160 X-ray instrument (Precision Instruments). Recipient mice were bled via the tail vein at the indicated intervals post-transplantation for analysis of peripheral blood donor chimerism. APC-labeled spherobeads were added to whole peripheral blood prior to staining with B220-APCy7, CD3-A700, Mac1-PECy7, Ter119-PECy5, Gr1-PB, CD71-Biotin/STA-BV605, and CD61-Alexa-647 (Biolegend) to detect mature lineage subsets for both host and donor mice. Cell suspensions were analyzed by FACS Aria III or LSR II (Becton Dickinson, San Jose, CA) for whole blood phenotypes and again post red blood cell lysis with a hypotonic alkaline lysis solution (ACK). Cells counts were calculated based on the number of cells analyzed and known number spherobeads added per sample.

#### *In vitro* Proliferation

Megakaryocyte progenitors (MkP: Lin^-^/cKit^+^/Sca1^-^/Slam^+^/CD41^+^) were prospectively isolated using CD117-Microbead enrichment (Miltenyi) and sorted by fluorescence-activated cell sorting (FACS) from young (6-12 weeks old) and old WT (20-24 months old) mice as previously described (Smith-Berdan *et al.*, 2015). 2000 cells were plated per well into a 96-well U-bottom tissue culture plate. Sorted cells were cultured for 3 days (MkP) in 200μl/well containing IMDM medium (Fisher) supplemented with 10% FBS, 50ng/ml rmTPO, 20ng/ml rmIL-6, 20ng/ml of rmIL-11, 50ng/ml of rmSCF, and 10ng/ml rmIL-3 (Cytokines from Peprotech) and Primocin (Invivogen). On days 3 APC-labeled spherobeads (BD Bioscience) were added to appropriate wells with cells and triplicate samples were analyzed using a FACSAria (Becton Dickinson, San Jose, CA). Cell expansion rates were calculated based on the number of beads recovered per beads added per well.

#### RNA-Seq

The RNA-Seq libraries were generated by sorting MkPs from young or old C57Bl6 mice using FACS Aria III (Becton Dickinson, San Jose, CA) as previously described. Cells were then stored in Trizol (Invitrogen) at −80°C until RNA isolation. RNA-Seq libraries were generated using Nextera Library Prep, as we have previously done (Beaudin *et al.*, 2016; Byrne *et al.*, 2017). Libraries were validated using the Bioanalyzer (Agilent 2100), validated RNA-Seq libraries were sequenced using Illumina HiSeq 4000 as Paired-end run reads at the QB3-Berkeley Genomics at University of California Berkeley and DESeq analysis was done with the help of Dr. Sol Katzman at the UCSC Bioinformatics Core. Heatmap was generated using Morpheus, https://software.broadinstitute.org/morpheus. Go Term analysis was run using Panther.

The datasets generated in the current study are available in the Gene Expression Omnibus (GEO), accession number GSEXXXX.

#### Quantification and Statistical Analysis

Number of experiments, n, and what n represents can be found in the legend for each figure. Statistical significance was determined by two-tailed unpaired student’s T-test. All data are shown as mean ± standard error of the mean (SEM) representing at least three independent experiments.

